# Lin28B is involved in curcumin-reversed paclitaxel chemoresistance and associated with poor prognosis in hepatocellular carcinoma

**DOI:** 10.1101/478479

**Authors:** Nan Tian, Wenbing Shangguan, Zuolin Zhou, Yao yao, Chunlei Fan, Lijun Cai

## Abstract

Chemoresistance remains a big challenge in hepatocellular carcinoma (HCC) treatment. Several studies indicated that RNA-binding protein Lin28B serves an oncogenic role in HCC, but its activity in HCC chemotherapy has never been assessed. In this study, we found that overexpression of Lin28B significantly increased the paclitaxel chemoresistance in two different HCC cells lines while silencing Lin28B reduced the chemoresistance in paclitaxel-resistance HCC cells. Curcumin, a natural anti-cancer agent, increased the sensitivity of HCC cells to paclitaxel through inhibiting NF-κB stimulated Lin28B expression both in vitro and in vivo. Furthermore, by analyzing TCGA (The Cancer Genome Atlas) LIHC (liver hepatocellular carcinoma) and GSE14520 databases, we found that Lin28B was highly up-regulated in HCC tissue compared with that in normal tissue and associated with α-fetoprotein levels, and that patients with Lin28B higher expression had a significant shorter overall survival time than those with Lin28B lower expression. Our data reveal that Lin28B may serve as a predictive biomarker and a treatment target to reverse HCC chemotherapy resistance in future clinical practice.

**Summary statement:** upregulation of Lin28B not only confers poor prognosis in HCC patients but also increases chemoresistance in HCC cells. Thus, Lin28B may serve as a predictive biomarker for use to reverse chemoresistance in clinical practice.

## Introduction

Hepatocellular carcinoma (HCC) is one of the most common solid-organ malignancies worldwide and it is particularly prevalent in China. Approximately 600,000 new cases of HCC are diagnosed every year, and 43% of them occur in China (Forner et al., 2012; Chen et al., 2016). Because of the insidious onset of HCC and the lack of symptoms in the early stage of the disease, only 10 to 20% of HCC tumors can be surgically excised. Therefore, for most cases of HCC, chemotherapy is the main treatment strategy (Nagano, 2010). However, hepatoma cells are known to be highly resistant to chemotherapeutic drugs, including paclitaxel (Wu et al., 2018; Lin et al., 2018a; Huang et al., 2018; Li et al., 2018). Chemoresistance has become a major obstacle in the treatment of HCC, clarifying the molecular mechanisms of drug resistance in HCC is important for developing more effective therapeutic approaches.

Mammalian genomes encode two evolutionarily conserved and developmentally regulated RNA-binding proteins, Lin28A and Lin28B, which are master regulators of the *let-7* family of microRNAs (Ambros and Horvitz, 1984; Moss et al., 1997). Lin28A and Lin28B regulate *let-7* microRNAs by binding to the terminal loop of the miRNA precursors and blocking their biogenesis, resulting in derepression of *let-7* targets, such as cell cycle proteins, cyclin-dependent kinases, growth factors, and ribosomal proteins (Wang et al., 2015). The expression levels of these two Lin28 homologues are significantly downregulated during cellular differentiation, and they are thought to be involved in tumorigenesis. They may also regulate the pluripotency of embryonic stem (ES) cells (Lin et al., 2018b; Xu et al., 2009; Darr and Benvenisty, 2009), and combined with pluripotency factors such as octamer-binding transcription factor 4 (Oct4), Nanog, sex determining region Y box 2 (Sox2), Lin28A can assist in reprogramming somatic cells to pluripotent stem cells (Darr and Benvenisty, 2009; Yu et al., 2007).

Lin28B was first cloned from and was shown to be overexpressed in human HCC cells and clinical samples (Guo et al., 2006), and it functions as an oncogene by promoting malignant transformation (Kugel et al., 2016; Schnepp et al., 2015), facilitating tumor-associated inflammation (Beachy et al., 2012; Iliopoulos et al., 2009), reprogramming metabolism, acquiring immortality, and evading immune destruction (Ma et al., 2018). Mice overexpressing *Lin28B* develop multiple tumors, including lymphoma, neuroblastoma, colonic adenocarcinoma, Wilms’ tumor, hepatoblastoma, and HCC (Beachy et al., 2012; Molenaar et al., 2012; Madison et al., 2013; Urbach et al., 2014; Nguyen et al., 2014), suggesting that *Lin28B* alone is sufficient to drive cancer. Furthermore, clinical epidemiological studies have indicated that Lin28B is associated with both susceptibility to certain cancers and overall survival in cancer patients (Permuth-Wey et al., 2011; Diskin et al., 2012; Chen et al., 2011). Lin28B is involved in the maintenance of liver cancer through several ways like downregulating the expression of anti-onco-miRNAs (Panella et al., 2018; Liang et al., 2010) and promoting Sp-1/c-Myc (You et al., 2014). However, the impact of Lin28B in HCC chemotherapy is unknown.

Curcumin, the golden nutraceutical derived from the roots of *Curcuma longa* Linn., has been reported to possess anti-cancer property against several tumor types, including HCC (Marquardt et al., 2015; Hu et al., 2018). It has been used to suppress inflammation, induce apoptosis, reduce tumor invasion and angiogenesis, and reverse drug resistance. Our previous experiments showed that curcumin sensitized HCC cells to the therapeutic agent paclitaxel. Therefore, in the current study, we generated a Lin28B-overexpresison cell line, a paclitaxel-resistant HCC cell line and a xenograft tumor model to explore whether Lin28B contributes to chemoresistance in HCC, and explore the underlying mechanism by which curcumin reverses paclitaxel resistance in HCC cells. Moreover, we used two independent Liver Hepatocellular Carcinoma gene expression datasets to evaluate the potential clinical value of Lin28B in HCC.

## Results

### Increased Lin28B mRNA expression confers poor prognosis in HCC patients

To investigate Lin28B mRNA expression in HCC, two independent liver hepatocellular carcinoma gene expression datasets (TCGA-LIHC and GSE41804) were employed. As shown in Fig. 1A and 1B, HCC tissues showed significant higher *Lin28B* transcript levels than normal tissues in both datasets (*p* <0.05). Next, we investigated the correlation between Lin28B expression and overall survival (OS) or progression-free survival (PFS) of HCC patients using Kaplan-Meier survival curve with log-rank tests. The analysis of the TCGA-LIHC cohort data revealed that patients with high Lin28B expression in tumor tissues exhibited significantly shorter survival than patients with low Lin28B expression levels (*p* =0.046; Fig. 1C). However, analysis of the TCGA-LIHC cohort revealed no correlation between Lin28B expression and PFS (Fig. 1D). These data indicate the potential clinical value of Lin28B in HCC.

**Figure 1.**
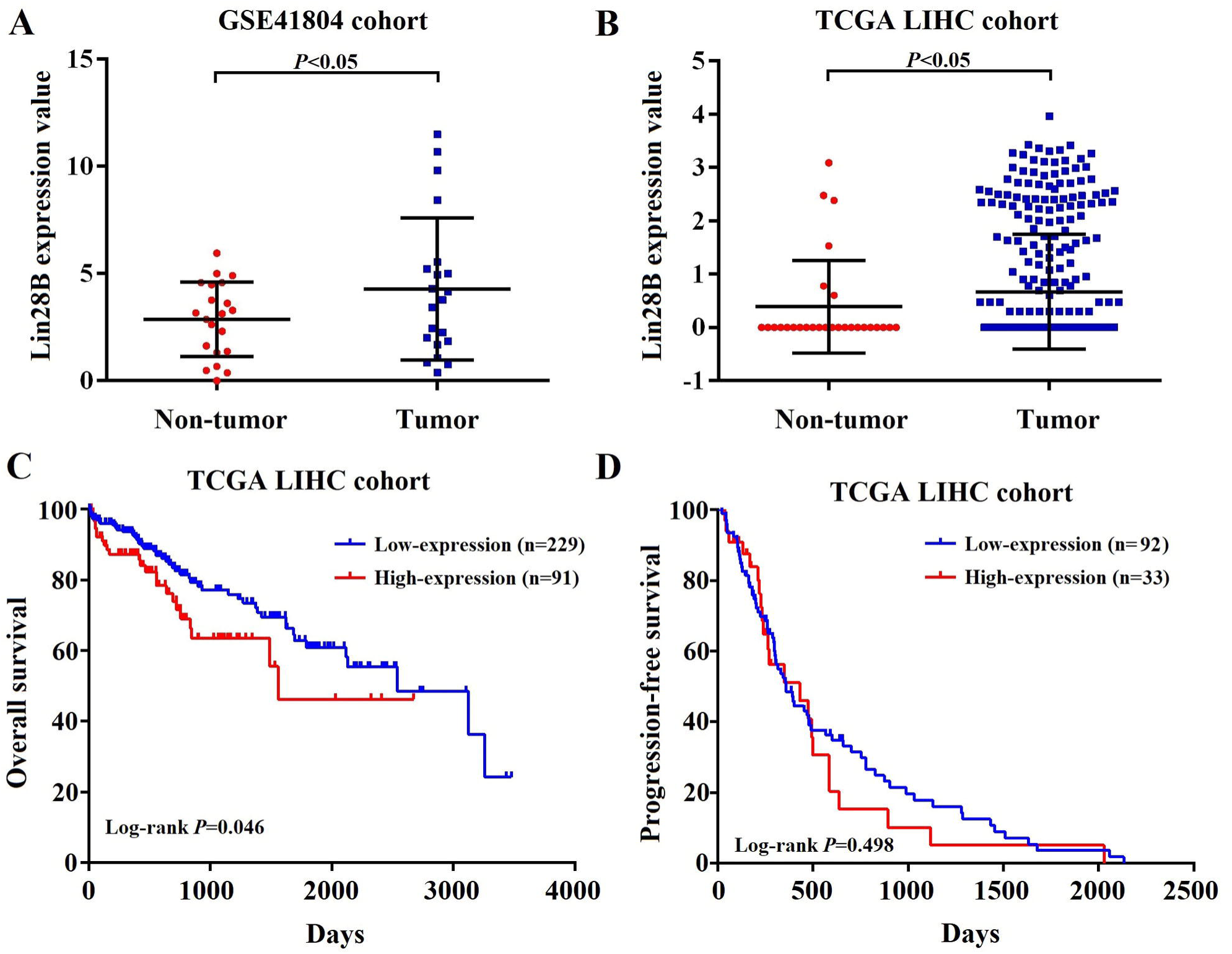
Increased Lin28B mRNA expression confers poor prognosis in patients with HCC. (A, B) Lin28B levels in the HCC tissues of patients in the GSE41804 and TCGA-LIHC datasets, respectively. Kaplan-Meier survival curve showing the (C) OS and (D) PFS rates in the TCGA-LIHC cohort. \**p* <0.05.

### Lin28B may be a valuable prognostic marker for patients with HCC

We evaluated the association between Lin28B expression and the clinicopathological features of patients with HCC by the Chi-square test to explore the potential oncogenic role of Lin28B in HCC. Statistical analysis of the TCGA-LIHC cohort indicated that Lin28B expression was positively associated with α-fetoprotein (AFP) levels (*p* =0.023). However, no statistically significant correlation was observed between Lin28B and other tested clinicopathological features, such as sex, age, neoplasm grade, pathologic stage, and relapse (Table 1). Next, univariate and multivariate analyses were performed to estimate the prognostic variables of Lin28B in the patients included in the TCGA-LIHC. Univariate analysis indicated that Lin28B expression was significantly associated with OS (hazard ratio [HR], 1.377; *p* = 0.029; 95% confidence interval [95% CI], 1.033-1.826; Table 2). Multivariate analysis showed that Lin28B might be an independent prognostic factor for patients with HCC (HR, 1.002; 95% CI, 1.001-1.002; *p* <0.001). These preliminarily results indicate that Lin28B may have potential clinical value as a predictive biomarker for OS in patients with HCC.

**Table I:** Correlation between Lin28B expression and HCC clinicopathological features in TCGA-LIHC

| Variable | Total, n | Lin28B expression, n | | $\chi^2$ | P-value |
| --- | --- | --- | --- | --- | --- |
|  |  | Low | High |  |  |
| Sex | 343 |  |  | 0.022 | 0.882 |
| Male | 234 | 165 | 69 |  |  |
| Female | 109 | 76 | 33 |  |  |
| Age (years) | 342 |  |  | 2.84 | 0.092 |
| $\leq 50$ | 74 | 58 | 16 | | |
| $> 50$ | 268 | 183 | 85 | | |
| Family cancer history | 295 |  |  | 0.463 | 0.496 |
| Yes | 94 | 70 | 24 |  |  |
| No | 201 | 142 | 59 |  |  |
| Lymphocyte infiltration | 214 |  |  | 0.785 | 0.675 |
| None | 104 | 72 | 32 |  |  |
| Mild | 93 | 69 | 24 |  |  |
| Severe | 17 | 13 | 4 |  |  |
| Vascular tumor cell invasion | 291 |  |  | 2.704 | 0.259 |
| None | 190 | 139 | 51 |  |  |
| Micro | 86 | 62 | 24 |  |  |
| Macro | 15 | 8 | 7 |  |  |
| Neoplasm grade | 340 |  |  | 1.059 | 0.303 |
| G1+G2 | 214 | 154 | 60 |  |  |
| G3+G4 | 126 | 84 | 42 |  |  |
| Pathologic stage | 322 |  |  | 0.716 | 0.398 |
| I+II | 240 | 170 | 70 |  |  |
| III+IV | 82 | 34 | 28 |  |  |
| AFP level (ng/ml) | 257 |  |  | 5.175 | 0.023* |
| $< 400$ | 197 | 148 | 49 | | |
| $\geq 400$ | 60 | 36 | 24 | | |
| Child-Pugh classification grade | 225 |  |  | 2.765 | 0.096 |
| A | 206 | 148 | 58 |  |  |
| B+C | 19 | 17 | 2 |  |  |
| Liver fibrosis Ishak score | 187 |  |  | 0.027 | 0.871 |
| $\leq 4$ | 114 | 84 | 30 | | |
| $> 4$ | 73 | 53 | 20 | | |
| Relapse | 258 |  |  | 0.809 | 0.368 |
| Yes | 91 | 67 | 24 |  |  |
| No | 167 | 114 | 53 |  |  |

\* $P < 0.05$ . AFP: α-fetoprotein.

**Table II:** Univariate and multivariate analysis of prognostic factors in patients included in the TCGA-LIHC

| Variable | Univariate analysis |  |  | Multivariate analyses |  |  |
| --- | --- | --- | --- | --- | --- | --- |
|  | HR | 95% CI | P-value | HR | 95% CI | P-value |
| Sex | 1.520 | 0.891-2.591 | 0.124 |  |  |  |
| Age (years) | 1.456 | 0.818-2.592 | 0.202 |  |  |  |
| Family cancer history | 0.739 | 0.431-1.266 | 0.270 |  |  |  |
| Lymphocyte infiltration | 0.813 | 0.569-1.163 | 0.257 |  |  |  |
| Vascular tumor cell invasion | 1.294 | 0.831-2.015 | 0.254 |  |  |  |
| Neoplasm grade | 0.724 | 0.445-1.176 | 0.191 |  |  |  |
| Pathologic stage | 0.942 | 0.427-2.076 | 0.882 |  |  |  |
| AFP level (ng/ml) | 0.754 | 0.423-1.345 | 0.339 |  |  |  |
| Child-Pugh classification grade | 1.099 | 0.421-2.866 | 0.847 |  |  |  |
| Liver fibrosis Ishak score | 0.915 | 0.572-1.464 | 0.712 |  |  |  |
| Relapse | 0.619 | 0.385-0.996 | 0.048* | 0.613 | 0.381-0.986 | 0.044* |
| Lin28B expression | 1.377 | 1.033-1.826 | 0.029* | 1.002 | 1.001-1.002 | <0.001*** |

\* $P < 0.05$ , \*\* $P < 0.01$ , \*\*\* $P < 0.001$ . AFP: α-fetoprotein, HR: hazard ratio, CI: confidence interval.

### Lin28B contributes to paclitaxel resistance in HCC cells

To further assess the contribution of Lin28B to paclitaxel resistance, we first overexpressed Lin28B in Hep3B cells via stable transfection of pEnter-Lin28B and then measured the IC_50_ of paclitaxel. As shown in Fig.2A, the IC_50_ of paclitaxel was increased in Hep3B cells stably overexpressing Lin28B, and the drug resistance index (RI) was 61.7% higher in Lin28B-overexpression cells than in cells transfected with a control vector (*p* <0.01). To confirm this finding, we next investigated the effect of Lin28B knockdown on paclitaxel resistance in Hep3B/TAX cells following transient transfection of Lin28B siRNAs. The results showed that two siRNAs successfully knocked down Lin28B expression in Hep3B/TAX cells, and effectively reduced the paclitaxel IC_50_ (Fig. 2B). Accordingly, high Lin28B expression contributed to paclitaxel resistance in Hep3B/TAX cells as compared to the resistance of parental Hep3B cells.

**Figure 2.**
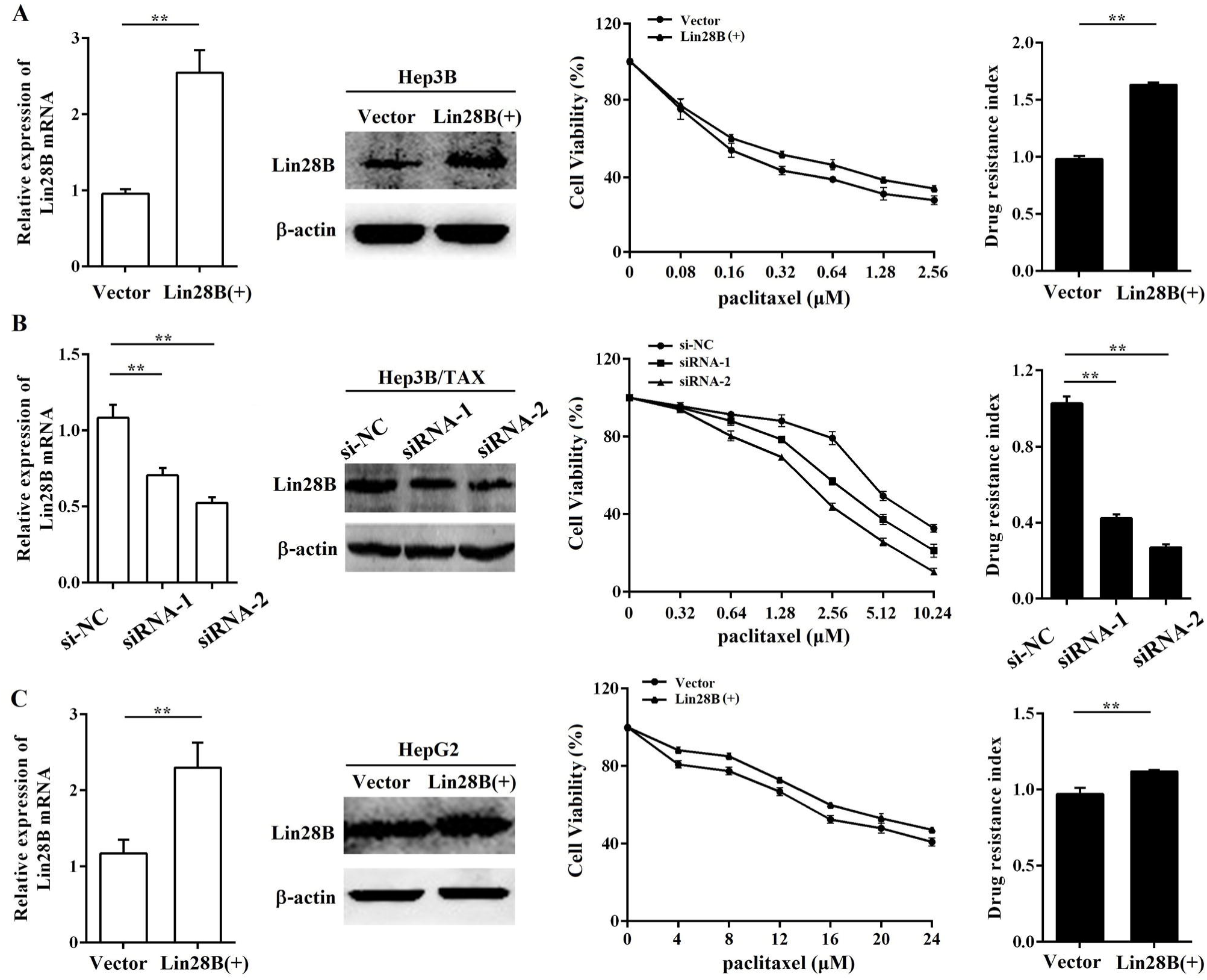
Contribution of Lin28B to paclitaxel resistance in HCC cells. (A) Hep3B cells stably overexpressing Lin28B, (B) Hep3B/TAX cells transfected with Lin28B siRNAs for 48h, and (C) HepG2 cells stably overexpressing Lin28B were subjected to qRT-PCR, western blot, and the MTT assay. Left, qRT-PCR and western blot to assess Lin28B expression; middle, representative dose-dependent cell viability curves; right, average resistance indexes (RI, the ratio of IC_50_ value of the experimental cells to that of the control cells). Vector, empty vector transfected; Lin28B (+), Lin28B overexpression; si-NC, negative control siRNA; siRNA, Lin28B siRNA; \**p* <0.05; \*\**p* <0.01.

To determine whether the contribution of Lin28B to paclitaxel resistance is specific to the Hep3B cell line, we investigated the effect of Lin28B on paclitaxel resistance in another HCC cell line, HepG2. We overexpressed Lin28B in HepG2 cells and determined the IC_50_ of paclitaxel by the MTT assay. The results showed that increased Lin28B expression significantly reduced the sensitivity of HepG2 cells to paclitaxel (Fig. 2C). Compared to the RI in cells transfected with a control vector, the RI in Lin28B-overexpression HepG2 cells was 11.2% higher (*p* <0.01). These results suggest that the expression of Lin28B contributes to paclitaxel resistance in HCC cells and that this effect is not specific to the Hep3B cell line.

### Lin28B suppresses paclitaxel-induced apoptosis in HCC cells

Paclitaxel has been shown to induce apoptosis in several cancer cells. Thus, we determined the effect of Lin28B on paclitaxel-induced apoptosis in HCC cells. Because Lin28B siRNA-2 was more efficient, we evaluated the apoptotic rate in Lin28B siRNA-2-transfected Hep3B/TAX cells treated with paclitaxel. Flowcytometry analysis showed that approximately 36.59 ± 6.63% of the Lin28B-silenced cells underwent apoptosis after treatment with 0.1μM paclitaxel, which was significantly higher than that in the control and si-NC cells (8.33 ± 0.56% and 8.63 ± 0.96%, respectively; Fig. 3A). In contrast, Hep3B and HepG2 cells, stably overexpressing Lin28B, were more resistant to paclitaxel-induced apoptosis, and Lin28B overexpression significantly reduced the number of apoptotic cells after paclitaxel treatment (*p*<0.01, Fig.3B) compared to the empty-vector transfected cells. Furthermore, the activities of apoptosis-related proteins, which are commonly used as markers of apoptosis, were also detected. As shown in Fig.3C, Lin28B knockdown increased the cleavage of caspase 9 and 3 and reduced the expression of the anti-apoptotic protein Bcl-2 in Hep3B/TAX cells treated with 0.1μM paclitaxel, whereas Lin28B overexpression reduced the cleavage of caspase 9 and 3 and increased the expression of Bcl-2 in Hep3B and HepG2 cells, as compared to the levels in their respective control cells.

**Figure 3.**
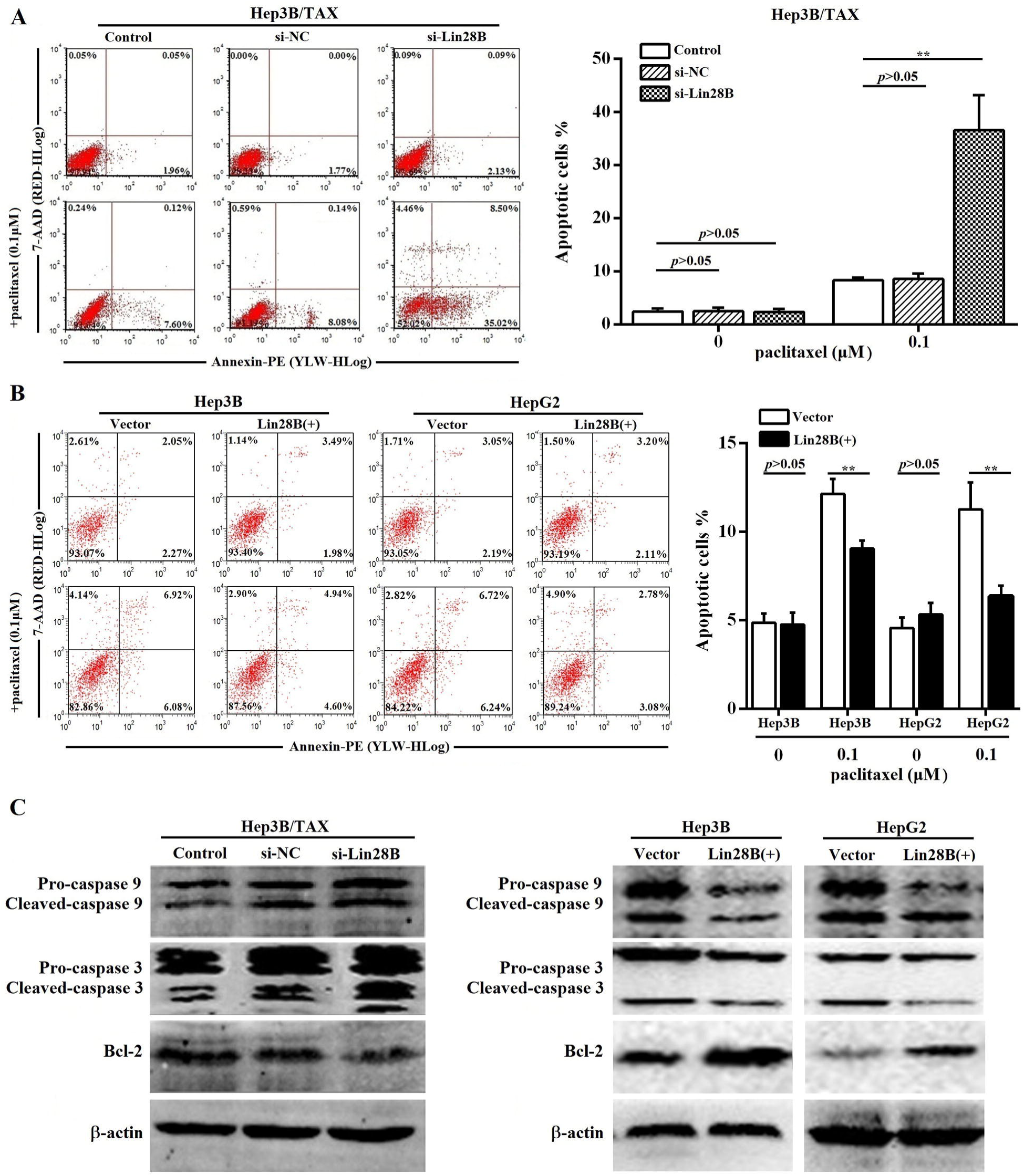
Effect of Lin28B on paclitaxel-induced apoptosis. (A) Hep3B/TAX cells transiently transfected with siRNAs for 48h and (B) Hep3B and HepG2 cells stably overexpressing Lin28B were treated with paclitaxel for 48 h and apoptosis was analyzed. (C) Western blot of apoptosis-related proteins (caspase 9, caspase 3, and Bcl-2). Vector, empty vector transfected; Lin28B (+), Lin28B overexpression; si-NC, negative control siRNA; si-Lin28B, Lin28B siRNA; \**p*<0.05; \*\**p*<0.01.

### Curcumin enhances paclitaxel-induced cytotoxicity and apoptosis in Hep3B/TAX cells

The results of the MTT assay showed that curcumin suppressed Hep3B/TAX cells proliferation with an IC_50_ of 34.99 ± 2.43 μM (Fig. 4A, IC_50_ of paclitaxel is 5.65 ± 0.51 μM). Therefore, we examined the effect of curcumin combined with paclitaxel at a series of concentrations lower than the IC_50_ to determine whether there is synergy between curcumin and paclitaxel. When combined with curcumin, the IC_50_ of paclitaxel in Hep3B/TAX cells was significantly decreased, in a dose-dependent manner. As shown in Fig.4B, when the concentration of curcumin was increased from 5 to 20 μM, the IC_50_ of paclitaxel decreased from 2.88 to 1.33 μM. Then, isobologram analysis was performed to evaluate the interaction between paclitaxel and curcumin. We found that the CI was considerably less than 1 when paclitaxel combined with curcumin, the CI was 0.65 for 2.88 μM paclitaxel and 5 μM curcumin, 0.66 for 2.09 μM paclitaxel and 10 μM curcumin, and 0.81 for 1.33 μM paclitaxel and 20 μM curcumin (Fig.4C). These results revealed that pretreatment of Hep3B/TAX cells with curcumin increased their sensitivity to paclitaxel.

**Figure 4.**
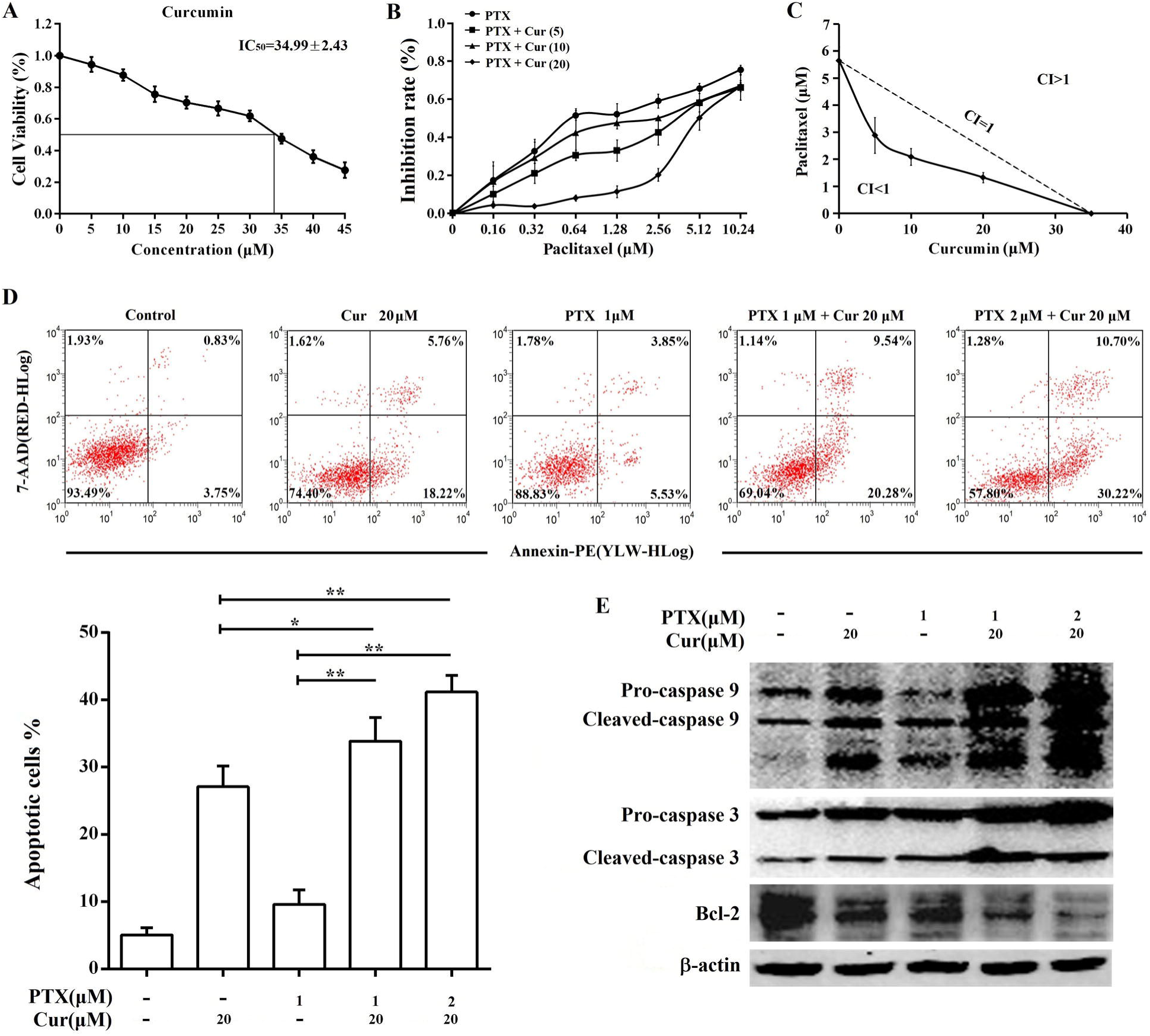
Co-treatment with curcumin and paclitaxel causes cytotoxicity and apoptosis in Hep3B/TAX cells. (A) MTT assay of Hep3B/TAX cells treated with increasing concentrations of curcumin for 48 h. The IC_50_ value was defined as the dose of the drug required to inhibit cell growth by 50% and was calculated using the improved Karber’s method. (B) Hep3B/TAX cells were treated with 1 μM paclitaxel combined with increasing concentrations of curcumin for 48 h, and then cell viability was analyzed by the MTT assay. (C) Isobologram analysis showing the effects of paclitaxel treatment combined with curcumin on Hep3B/TAX cells. Hep3B/TAX cells were stained with Annexin V-PE/7-AAD and analyzed by (D) flow cytometry and (E) western blot, after treatment with paclitaxel and curcumin individually, or in combination for 48 h. Cur, curcumin; PTX, paclitaxel; * *p*<0.05; ** *p*<0.01.

The ability of curcumin to enhance paclitaxel-induced apoptosis in Hep3B/TAX cells was assessed by an apoptosis assay. Cells were treated with curcumin (20 μM), paclitaxel (1 μM), or co-treated with curcumin and paclitaxel for 48 h, and then an Annexin V PE/7-AAD apoptosis assay was performed to determine the population of apoptotic cells. It was observed that the combination of paclitaxel and curcumin resulted in significantly higher cell death than single agent chemotherapy (Fig. 4D). While individual treatment with paclitaxel and curcumin led to 10% and 27% cell death, respectively, the combination of 1 μM paclitaxel and 20 μM curcumin resulted in nearly 34% celldeath, and 2 μM paclitaxel and 20 μM curcumin resulted in 41% cell death. Moreover, paclitaxel-induced cleavage of caspase 9 and 3 was also increased by curcumin treatment, as shown in Fig.4E. Although individual treatment with 20 μM curcumin or 1 μM paclitaxel induced activation of caspase 9 and 3, combination chemotherapy increased the level of cleaved caspase in a dose-dependent manner. Taken together, these results indicate that curcumin sensitizes Hep3B/TAX cells to paclitaxel *in vitro*.

### Curcumin enhances paclitaxel-induced cytotoxicity and apoptosis in xenograft tumors

Next, we sought to determine whether curcumin sensitizes HCC cells to paclitaxel *in vivo*. A subcutaneous xenograft model in nude mice was established using Hep3B/TAX cells, and then treated with curcumin (40 mg/kg), paclitaxel (10 mg/kg), or curcumin (40 mg/kg) and paclitaxel (10 mg/kg). The results demonstrated that the growth of tumor was significantly suppressed in mice co-treated with of curcumin and paclitaxel as compared to those in the control as well as in mice treated with either curcumin or paclitaxel alone (Fig. 5A and 5B). Tumors from co-treatment group showed decreased immunohistochemical (IHC) staining for Ki-67compared to the controls and individually treated tumors (Fig. 5D). Moreover, as shown in Fig.5C, the expression of apoptosis-related proteins was significantly reduced in tumors in the combination treatment group compared with the control and individually treated tumors. These results provide evidence that curcumin sensitized resistant HCC cells to paclitaxel *in vivo*.

**Figure 5.**
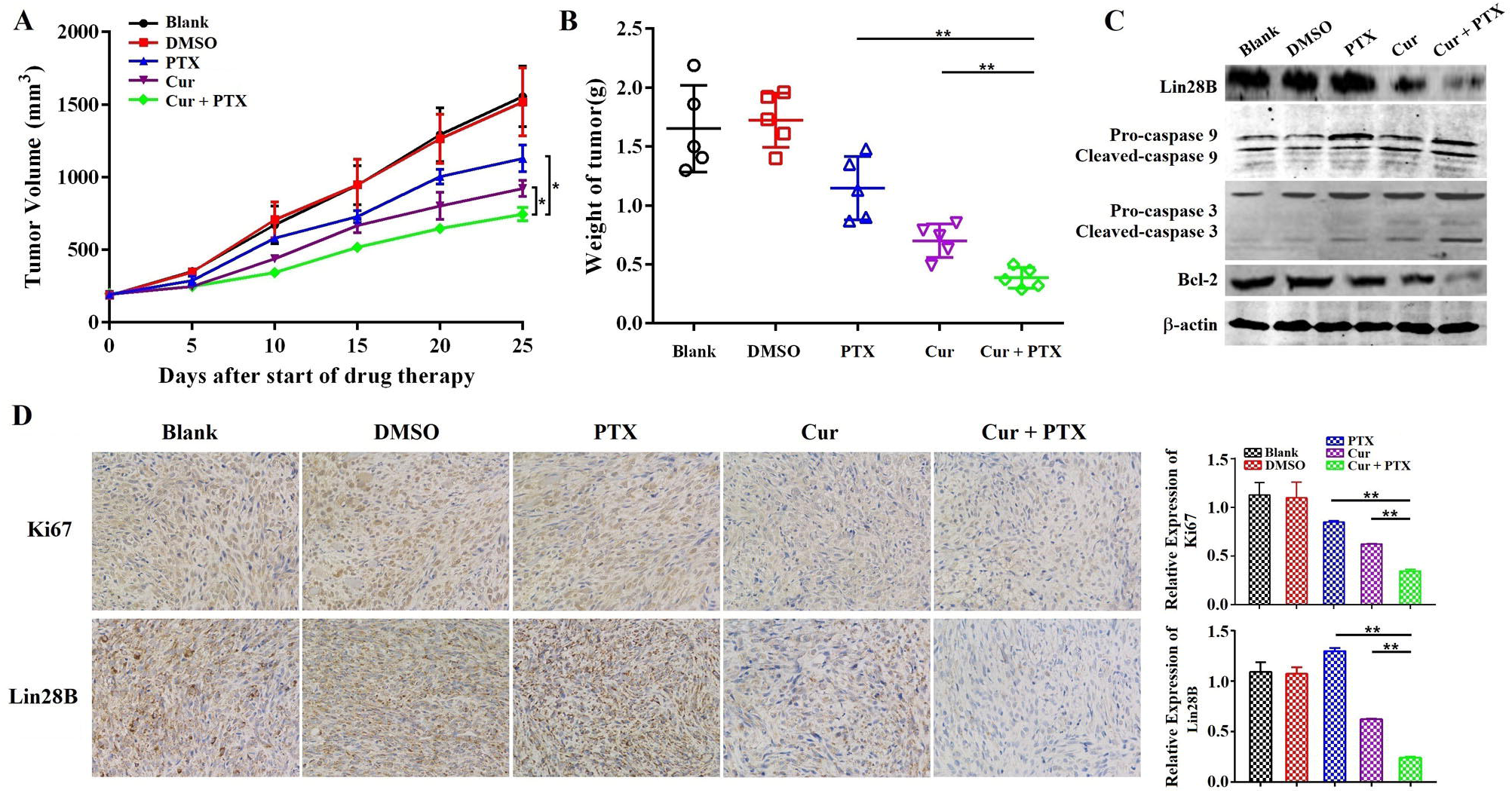
Co-treatment with curcumin and paclitaxel causes cytotoxicity and apoptosis in HCC cells *in vivo*. (A-B) Analysis of the anti-tumor effects of co-treatment with curcumin (40 mg/kg) and paclitaxel (10 mg/kg). Tumor volume is presented as a growth curve, and the tumor weight is presented as a Cleveland dot plot. (C) Western blot of apoptosis-related proteins (caspase 9, 3, and Bcl-2) and Lin28B expression levels in the tumors of the mice. (D) Representative immunohistochemistry (IHC) images of Ki-67 and Lin28B in tumor tissue samples. \**p*<0.05; \*\**p*<0.01.

### Curcumin sensitizes HCC to paclitaxel by downregulating Lin28B and NF-B

To clarify the molecular mechanism by which curcumin reverses paclitaxel resistance in HCC cells, we first evaluated the expression of Lin28B in paclitaxel treated, and curcumin and paclitaxel co-treated Hep3B/TAX cells. As shown in Fig.6A and 6B, Lin28B expression was increased upon stimulation with paclitaxel; however, this increase was blocked by curcumin in a dose-dependent manner. Moreover, combined treatment with curcumin and paclitaxel downregulated the expression of Lin28B in the xenograft tumors (Fig. 5C and 5D), suggesting that the paclitaxel-induced increase in Lin28B expression was inhibited by curcumin.

**Figure 6.**
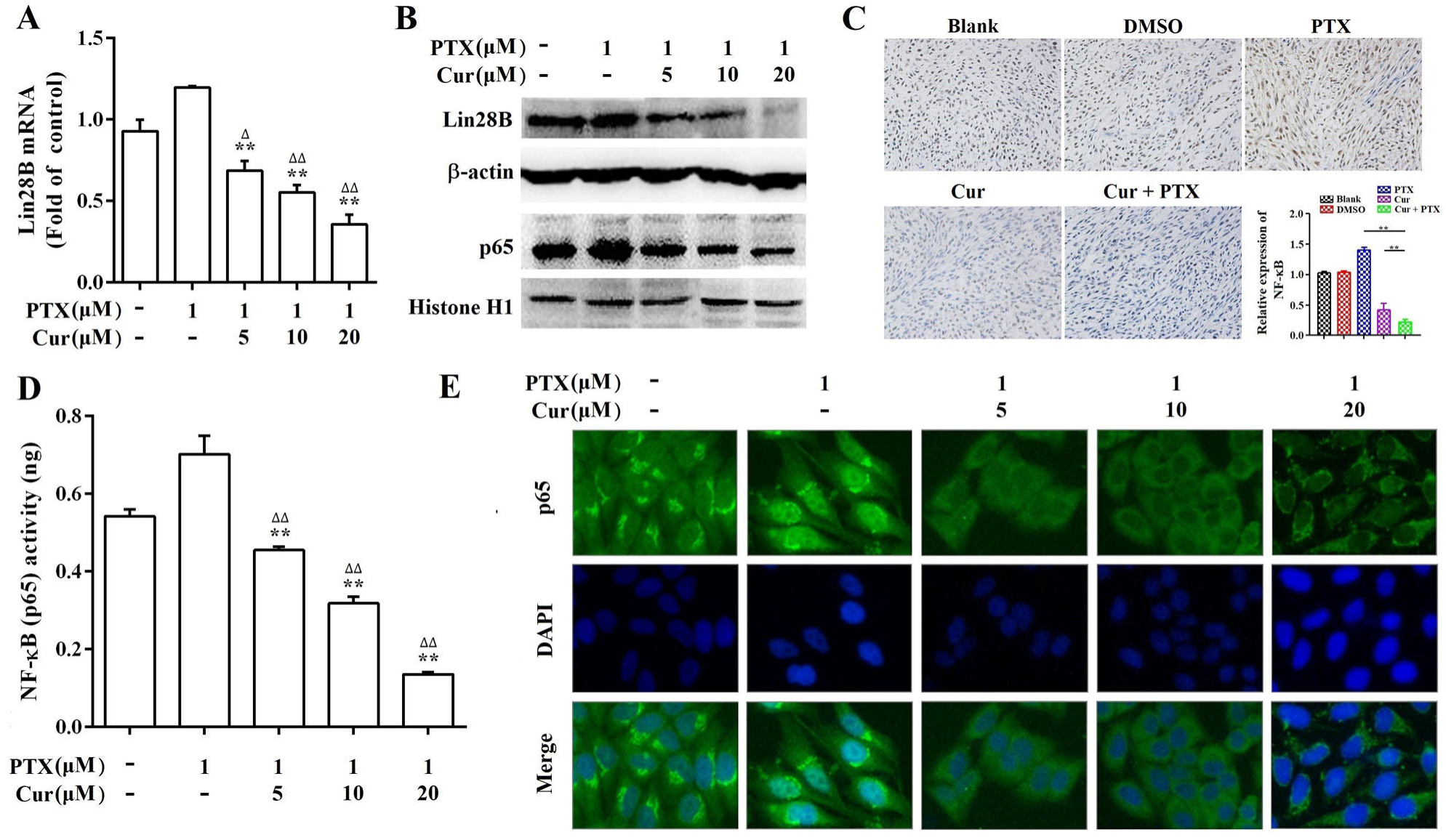
The effect of curcumin on paclitaxel-induced Lin28B and NF-κB levels. (A, B) qRT-PCR and western blot showing Lin28B levels in Hep3B/TAX cells treated with curcumin and paclitaxel alone or in combination.(C) IHC showing NF-κB levels in the nuclear fractions of Hep3B/TAX cells treated with curcumin and paclitaxel alone or in combination. (D) Hep3B/TAX cells were fixed with 4% paraformaldehyde and permeabilized with 5% Triton X-100. After blocking the slides with 5%BSA, they were stained for NF-κB and mounted using DAPI, and images were captured using a Nikon microscope. (E) NF-κB activity was measured using the NF-κB (p65) Transcription Factor Assay Kit. \**p*<0.05, ** *p*<0.01 for the control versus paclitaxel groups. ^∆^*p*<0.05 ^∆∆^*p*<0.01 for paclitaxel versus paclitaxel and curcumin co-treatment groups (Cur, curcumin; PTX, paclitaxel).

The inflammatory transcription factor NF-κB has been shown to directly activate Lin28B, and NF-κB is reported to be highly expressed and activated by paclitaxel in several tumor cell lines including Hep3B (Iliopoulos et al., 2009; Shuang et al., 2017). Therefore, we performed a series of experiments to analyze the expression, nuclear translocation, and DNA-binding capacity of NF-κB in Hep3B/TAX cells. Western blot and IHC staining revealed that co-treatment with curcumin and paclitaxel inhibited NF-κB protein expression more than paclitaxel treatment alone both *in vitro* and *in vivo* (Fig.6B and 6C). The results of immunofluorescence staining (IF) and NF-κB (p65) transcription factor assay showed a significant decrease in NF-κB translocation and dose-dependent suppression of NF-κB DNA binding in curcumin and paclitaxel co-treated Hep3B/TAX cells (Fig.6D and 6E).

### Stimulation of NF-κB abrogates curcumin-inhibited Lin28B expression

It is well known that tumor necrosis factor-alpha (TNF-α) is a classical activator of NF-κB signaling. Therefore, we stimulated Hep3B/TAX cells with 10 ng/mL TNF-α for 0.5, 1, and 2 h after a 48h curcumin treatment to evaluate whether curcumin inhibits the expression of Lin28B through NF-κB. As shown in Fig.7A, 7C, and 7D, 10 ng/mL TNF-α significantly increased the expression, DNA-binding, and translocation activities of NF-κB in cell nuclei, which was inhibited by a 48 h treatment with curcumin. Moreover, curcumin-inhibited Lin28B expression was stimulated by TNF-α in a time-dependent manner (Fig.7A and 7B), indicating that curcumin inhibited the expression of Lin28B through NF-κB.

**Figure 7.**
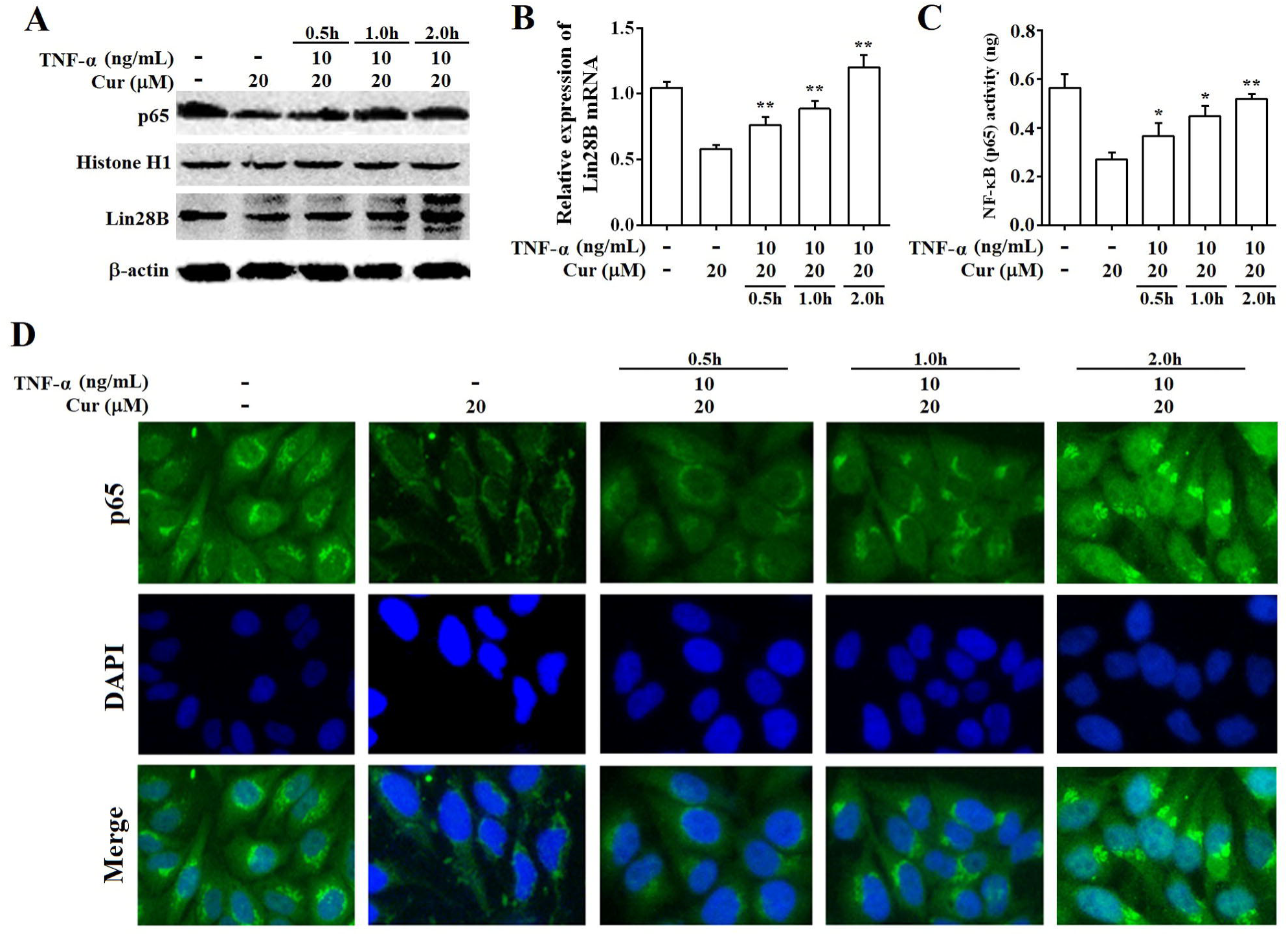
Stimulation of NF-κB abrogates curcumin-inhibited Lin28B expression. (A) Western bolt analysis of Lin28B and NF-κB (p65) expression levels in Hep3B/TAX cells, following stimulation with 10 ng/mL TNF-α and a 48 h curcumin treatment. (B) qRT-PCR analysis of Lin28B in Hep3B/TAX cells stimulated by 10 ng/mL TNF-α after a 48 h curcumin treatment. (C) DNA-binding activity of NF-κB (p65) was measured using the NF-κB (p65) Transcription Factor Assay Kit. (D) Immunofluorescence staining was performed, and images were captured using a Nikon microscope. \**p*<0.05 \*\**p*<0.01 for control versus paclitaxel groups. ^∆^*p*<0.05 ^∆∆^*p*<0.01 for paclitaxel versus paclitaxel and curcumin co-treatment groups (Cur, curcumin; PTX, paclitaxel).

## Discussion

Several studies have reported the aberrant expression of Lin28B and its association with outcomes in epithelial ovarian cancer, breast cancer, and neuroblastoma. In the present study, we investigated the correlation between Lin28B expression and clinical outcomes in HCC patients as well as its role in the susceptibility of HCC to chemotherapy. The results showed that Lin28B is overexpressed in HCC tissues and is closely associated with a patient’s AFP levels, which is used as a diagnostic criterion for HCC. In addition, univariate and multivariate analyses indicated that high Lin28B expression predicts poorer prognosis of HCC patients, and may act as a prognostic biomarker. Cheng *et al.* (Cheng et al., 2013) demonstrated that, for HCC, Lin28B is associated with a high tumor grade, large size, high American Joint Committee on Cancer (AJCC) stage, high Barcelona-Clinic Liver Cancer (BCLC) stage and recurrence. However, we did not observe a correlation between Lin28B expression and these clinicopathological features. This may be due to a difference in Lin28B assessment methods, as we measured Lin28B levels in the solid tumor, whereas Cheng *et al.* measured circulating Lin28B levels.

Chemoresistance is a major cause of treatment failure and is an important challenge for cancer therapy (Kim et al., 2017). Therefore, identifying the molecular markers of chemoresistance and increasing tumor-cell sensitivity to chemotherapeutic drugs are critical for enhancing therapeutic efficacy. Hsu *et al.* (Hsu et al., 2015) demonstrated that overexpression of Lin28B was associated with resistance to platinum-based chemotherapy in ovarian cancer. In this study, we found that overexpression of Lin28B significantly increased paclitaxel-induced chemoresistance in Hep3B and HepG2 cells lines, while silencing of Lin28B reduced chemoresistance in Hep3B/TAX cells, indicating that Lin28B contributes to paclitaxel-induced chemoresistance in HCC cells. Paclitaxel has been used to treat a wide range of tumors; its anti-tumor mechanism involves inhibition of microtubule assembly, which blocks the progression of mitosis and prolonged activation of this mitotic checkpoint triggers apoptosis (Bharadwaj and Yu, 2004; Brito et al., 2008). Our results showed that Lin28B inhibited paclitaxel-induced apoptosis in HCC cells and this effect may be related to the regulation of apoptosis-related protein activity. Thus, we speculate that Lin28B may be a useful molecular marker of chemoresistance in HCC.

An effort that deserves attention is a novel combination treatment strategy, that aims to reverse chemoresistance through the sequential addition of an adjuvant drug that targets resistance caused by chemotherapy (Yap et al., 2013). In our previous study, curcumin was found to sensitize HCC cells to paclitaxel *in vitro* (Zhou et al., 2015). In the current study, we use the paclitaxel-resistant cell line Hep3B/TAX and a xenograft mouse model to further explore the underlying mechanism of curcumin reversed paclitaxel-induced chemoresistance in HCC. Isobolograms showed a dose-dependent cytotoxic effect of paclitaxel on Hep3B/TAX cells, which was synergistically potentiated by co-treatment with curcumin. An annexin V PE/7-AAD apoptosis assay revealed that induction of apoptosis by paclitaxel in Hep3B/TAX cells was significantly enhanced by curcumin. Similarly, in the Hep3B/TAX-derived xenograft mouse model, we showed that co-treatment with curcumin and paclitaxel not only significantly reduced tumor size but also increased tumor cell apoptosis. Collectively, our findings, from both the *in vitro* and *in vivo* assays, provide valuable insights into the beneficial effects of curcumin on paclitaxel-induced chemoresistance in HCC.

Several studies have shown curcumin to be a promising sensitizer to chemotherapeutic agents, including paclitaxel, in a wide variety of tumor cell types (Kumar et al., 2017; Zhang et al., 2017; Zou et al., 2018; Dang et al., 2015; Lu et al., 2017; Zhan et al., 2014). Curcumin combined with standard docetaxel chemotherapy led to a significant improvement and clinical response in patients with metastatic breast cancer in a phase I clinical trial (Bayet-Robert et al., 2010). Considering the pleiotropic activity of curcumin in cancer prevention, it could be involved in chemosensitization in several different ways, such as inhibition of NF-κB signaling (Dang et al., 2015), miRNA-induced suppression of cell proliferation (Lu et al., 2017), and modulation of epidermal growth factor receptor (EGFR) signaling (Zhan et al., 2014). Our results showed that Lin28B expression in Hep3B/TAX cells and xenograft tumor tissue was significantly decreased in a dose-dependent manner following co-treatment with curcumin and paclitaxel. These results indicate that curcumin exhibits its paclitaxel-sensitizing effect by reducing of Lin28B expression in HCC.

Experiments involving sequence analysis combined with chromatin immunoprecipitation assay conducted in Kevin Struhl’s laboratory (Iliopoulos et al., 2009) revealed a highly conserved NF-κB motif in the first intron of the *Lin28B* gene, indicating that NF-κB activates *Lin28B* expression through direct binding. NF-κB, which is an important transcription factor for immunity, inflammation, and cell survival and is a hallmark of tumorigenesis (Kang et al., 2016), typically exists as a heterodimer composed of the Rel-family proteins p50 and p65 (RelA). In cancer, NF-κB/p65 regulates the transcription of growth-promoting and anti-apoptotic genes. Given the fact that curcumin has a potential to disrupt several steps of the NF-κB activation pathway, we further investigated the function of NF-κB/p65 in curcumin-inhibited Lin28B expression. Using the NF-κB signaling activator TNF-α, we found that curcumin-inhibited Lin28B expression was stimulated by TNF-α in a time-dependent manner, indicating that curcumin inhibits the expression of Lin28B through NF-κB.

In a conclusion, our data demonstrate that Lin28B is upregulated in paclitaxel-resistant HCC cells, and Lin28B overexpression results in paclitaxel resistance, while silencing Lin28B increased the sensitivity of to paclitaxel. Curcumin sensitized cells to paclitaxel-induced chemoresistance by inhibiting Lin28B and the upstream protein NF-κB (Fig. 8). Moreover, upregulation of Lin28B is associated with poor prognosis in patients with HCC. Thus, Lin28B may serve as a predictive biomarker for use in the development of personalized treatment strategies and as a treatment target to reverse chemoresistance in clinical practice.

**Figure 8.**
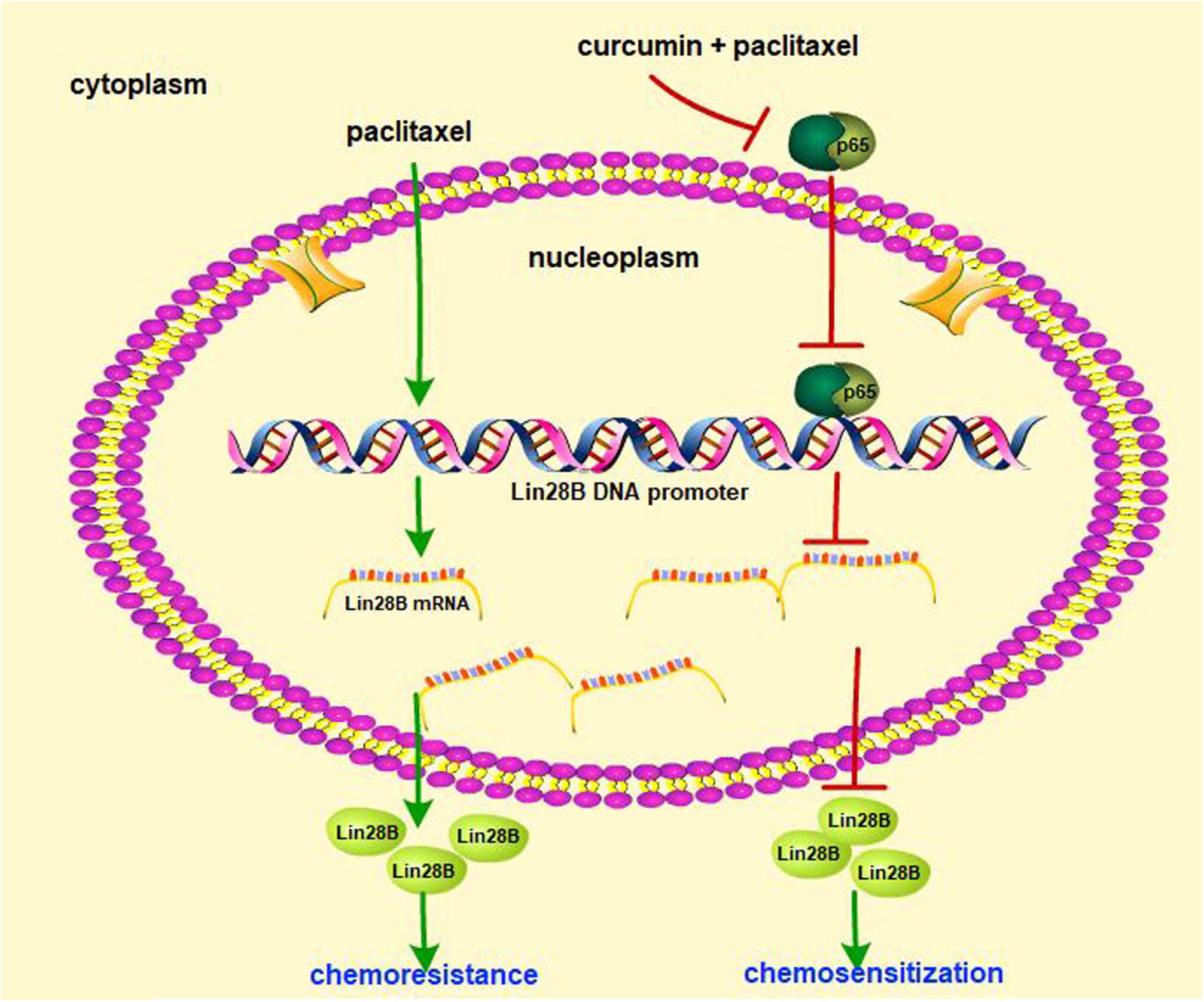
Schematic of paclitaxel-induced chemoresistance involving Lin28B and the chemosensitizing effect of curcumin in HCC cells. Paclitaxel treatment of HCC cells upregulated Lin28B expression and induced chemoresistance, whereas co-treatment with curcumin and paclitaxel downregulated Lin28B expression by inhibiting the activity of NF-κB.

## Materials and methods

### Cell lines and compounds

The human hepatoma cell line Hep3B, HepG2 were obtained from Cell Resource Center of Shanghai Institute of Life Sciences and grown in Dulbecco’s modified Eagle’s medium (DMEM) supplemented with 10% fetal calf serum (FBS), penicillin (100 U/mL) and streptomycin (100 mg/mL) at 37℃ in 5% CO_2_. Paclitaxel and curcumin were purchased from Tianjin Yi Fang Science and Technology, Ltd., (Tianjin, China), and dissolved in dimethyl sulfoxide (DMSO), stored at −20℃ and diluted in cell culture medium immediately prior to use.

### Human tissue samples

HCC gene expression datasets are deposited at NCBI’s Gene Expression Omnibus (GEO) public database (http://www.ncbi.nlm.nih.gov/geo/, GEO accession number, GSE41804) and The Cancer Genome Atlas Liver Hepatocellular Carcinoma (https://cancergenome.nih.gov/, TCGA-LIHC). GSE41804 contains 40 samples including 20 pairs of fresh HCC specimens and their corresponding adjacent non-tumor. TCGA-LIHC dataset contains 377 HCC cases and comprises follow-up information for patients and gene expression data. Overall survival (OS) and progression-free survival (PFS) were assessed and calculated from the follow-up data of the HCC patients in TCGA-LIHC. OS was defined as the period from the date of the pathological diagnosis to death; PFS was defined as the period from the date of the initial treatment to HCC progression, presenting as new tumor event.

### Development of a paclitaxel-resistant cell line (Hep3B/TAX)

To develop a paclitaxel-resistant HCC cell line, Hep3B cells were exposed to gradually increasing concentrations of paclitaxel (0.01-0.2 μM) in complete medium. Briefly, Hep3B cells were seeded in culture flasks at a density of 4~5 × 10 ^5^ cells/mL and allowed to grow. After 24 h incubation, paclitaxel (0.01 μM) was added, and the cells were incubated for another 24 h. Then, the cells were washed 3 times with D-Hanks solution and the medium was changed to paclitaxel-free medium. The cells were incubated and allowed to grow until confluent. Then, the cells were subcultured and re-exposed to double the dose of drug. This process was repeated until the cells were resistant to 0.2 μM paclitaxel. After successful development, Hep3B/TAX cells were maintained in complete medium containing a low concentration of paclitaxel (0.01 μM).

### Viability and apoptosis

The cell viability was measured by MTT assay according to the manufacturer’s protocol (Nanjing KeyGen Biotech. Co. Ltd., China). 5 × 10 ^3^ cells were plated on 96-well plates and after overnight incubation treated with the indicated drugs for 48 hours. MTT solution (5 mg/mL) was added to each well and the plate was incubated for 4 h at 37℃. After removal of the medium, formazan crystals were dissolved in 150 mL of DMSO. The absorbance of MTT-formazan was measured at 550 nm using a SpectraMaxM3 microplate reader (MolecularDevices, Sunnyvale, CA, USA). Apoptosis was assessed using Annexin V PE/7-AAD apoptosis assay (Nanjing KeyGen Biotech. Co. Ltd., China), stained cells were quantified using a Guava easyCyte 5 Flow Cytometer (EMD Millipore, USA).

### Isobologram analysis

For combination assay, IC_50_ was calculated from two sets of concentration response graphs, the combination index (CI) was calculated according to the following equation (Agarwal et al., 2015): Sum CI (ΣCI) =IC_50_ of A in mixture/ IC_50_ of A alone + IC_50_ of B in mixture/ IC_50_ of B alone. The CI values were defined as follows: <1, synergism; =1, additive; and >1, antagonism.

### RNA extraction and quantitative real-time PCR (qRT-PCR)

Total RNA from the cells was extracted using Trizol reagent (Thermo Fisher Scientific, USA). qRT-PCR was performed with a HiFi-MMLV cDNA kit and UltraSYBR Mixture (Beijing ComWin Biotech Co., Ltd., Beijing, China). The primers used for qRT-PCR to detect Lin28B (forward primer 5’-GACCCAAAGGGAAGACAC-3’, reverse primer 5’-TCTTCCCTGAGAACTCGCGG-3’) and β-actin (forward primer 5’-GGCACCACACCTTCTACAAT-3’, reverse primer 5’-GTGGTGGTGAAGCTGTAGCC-3’) were synthesized by Nanjing GenScript Co. Ltd., China. The fold-change of each gene was calculated using the 2^−ΔΔCt^ method.

### Preparation of nuclear extracts

Experiments were performed according to manufacturer’s instructions (nuclear extraction kit; KeyGen Biotech, Nanjing, China). Protein concentration was determined using a Nanodrop ND-1000 Spectrophotometer (NanoDrop Technologies, Wilmington, DE). The nuclear extract was used for western blot analysis.

### Western blot analysis

Whole cell lysates or tissues extracts were prepared using RIPA Lysis Buffer (Beyotime Biotechnology, China) containing PMSF Protease Inhibitor (Thermo Fisher Scientific, USA), separated by SDS-PAGE and transferred onto PVDF Membrane (Merck Millipore, Germany). Membranes were probed with the indicated antibodies. Mouse polyclonal anti-β-actin antibody was purchased from HuaBio Biotechnology (Huabio, China), Rabbits polyclonal antibodys of NF-κB, caspase 9, caspase 3, Bcl-2, Histone H1 were purchased from Proteintech Group, and anti-Lin28B rabbit polyclonal antibody was purchased from BBI Life Sciences Corporation. Immune complexes were detected using the enhanced chemiluminescence system (Advansta, Menlo Park, CA, USA).

### RNA interference

The short interfering RNA (siRNA) strand oligomers specific for Lin28B and its negative control siRNA (si-NC) were synthesized by Biomics Biotechnologies Co., Ltd. (Nantong, China). Cells were plated in 6-well plates and incubated for 24 h. Next, cells were transfected with siRNA using the Lipofectamin 2000 (Thermo Fisher Scientific, USA.) in accordance with the manufacturer’s instructions. At 24 h after transfection, cells treated with or without paclitaxel for another 48h, then harvested and used for MTT assay.

### Establishment of Lin28B stable expression cell lines

Cells were transfected with either pEnter or pEnter-Lin28B (Vigene Biosciences, China) using Lipofectamin 2000. After 72h incubation, puromycin was then added to the transfected cells for generating cells stably overexpressing Lin28B. Lin28B stable expression cell lines were identified by western blot, and then used for the experiments.

### Mouse tumor xenografts

BALB/c nude mice (male, 5-week-old, 16-20 g) were purchased from Shanghai Slack Laboratory Animal Co. Ltd. (animal license number: SCXK (Shanghai) 2012-0002). All procedures were performed with approval of and in accordance with the guidelines of the National Institutes of Health animal care committee. Hep3B/TAX cells were injected into the dorsal flanks of each mouse. After 14 days, the mice were randomly assigned to five groups (n=5), and either treated with or without DMSO, curcumin (40 mg/kg), paclitaxel (10 mg/kg), or curcumin plus paclitaxel by intraperitoneal injections every 3 days, respectively. Tumor volume was calculated using the formula V = W^2^ × L × 0.5. The mice were sacrificed and tumors were harvested for the follow -up experiments.

### Immunofluorescent staining (IF)

Cells grown on glass coverslips were treated with different concentrations of curcumin (0, 5, 10, and 20 μM) for 4 h and then co-treated with 1 μM paclitaxel for an additional 24 h. After washing with PBS, slides were fixed with 4% paraformaldehyde. Then after washing with 5% Triton-X100 for 15 min at 25℃, and slides were blocked with 5% BSA for 1 h and incubated with the NF-κB antibody overnight at 4℃. Following incubation with goat anti-rabbit secondary antibody at 25℃ for 2 h, slides were incubated with SYBR-FITC (Boster Bioengineering, Wuhan, China) for 1 h. Then DNA was counterstained with DAPI and images were captured with a Nikon ECLIPSE Ti-S microscope (Nikon, Japan).

### Immunohistochemistry (IHC)

The dissected tumors sections (5 μm) mounted on the glass slides were deparaffinized and rehydrated. Subsequently, the specimens were washed in 0.01 M PBS, treated with 3% hydrogen peroxide, and washed in 0.01 M PBS again. Blocking was performed with 3% BSA in 0.01 M PBS. Then the sections were incubated with antibodies for proliferation marker Ki-67 (Calbiochem, Cambridge, MA), Lin28B, and NF-κB, followed by incubation with secondary antibodies. After the cell nuclei were labeled with hematoxylin, images were captured using a microscope (XSP-C204, Chongqing Optical Instrument Co., Ltd.).

### NF-κB (p65) transcription factor assay

NF-κB activity level was measured by evaluating NF-κB (p65) DNA binding activity in nuclear extracts of Hep3B/TAX cells. Analysis was performed following the manufacturer’s protocol for NF-κB (p65) transcription Factor Assay Kit (Abnova, Germany). Cells were harvested and washed twice with PBS, 500 mL of cold hypotonic buffer supplemented with 0.5 mL of DTT, 5 mL of PMSF and 0.5 mL of protease inhibitor was added. Following centrifugation, the pellet containing the nuclear portion was then re-suspended in ice-cold complete nuclear extraction buffer and incubated on ice for 20 min. The supernatant fractions were collected and added to the plate with transcription factor binding assay buffer. Following incubation with primary and secondary antibodies, color development was detected after addition of the developing solution. Finally, sample absorbance was measured at 450 nm using a SpectraMaxM3 microplate reader (MolecularDevices, Sunnyvale, CA, USA).

### Statistical analysis

Statistical analysis was performed using Student’s *t*-test for pairwise comparison, 1-way ANOVA test for multiple group comparisons. *p*-values < 0.05 were considered statistically significant. Results were presented as means ± SD. Survival curves were constructed with the Kaplan-Meier method and analyzed using log-rank tests. The correlation between Lin28B expression and HCC clinicopathological features was analyzed using Pearson’s chi-square test. Prognostic factors were examined with univariate and multivariate Cox regression models.

## Acknowledgements

The authors would like to thank Dr Ting Zhang (Medical technology College, Zhejiang Chinese Medical University) for critical reading of the manuscript.

## Authors’ contributions

NT, CLF, and LJC conceived and designed the experiments. NT, SGWB, ZZL, and YY performed the experiments. SGWB, ZZL, and YY analyzed clinical data. NT, SGWB, and LJC performed data analysis. TN wrote the paper. All authors performed critical review of the manuscript. All authors read and approved the final manuscript.

## Consent for publication

Not applicable.

